# Subsensory stochastic electrical stimulation targeting muscle afferents alters gait control during locomotor adaptations to haptic perturbations

**DOI:** 10.1101/2022.11.08.515667

**Authors:** Giacomo Severini, Alexander Koenig, Iahn Cajigas, Nicholas Lesniewski-Laas, Jim Niemi, Paolo Bonato

## Abstract

Subsensory noise stimulation targeting sensory receptors has been shown to improve balance control in healthy and impaired individuals. However, the potential for application of this technique in other contexts is still unknown. Gait control and adaptation rely heavily on the input from proprioceptive organs in the muscles and joints. Here we investigated the use of subsensory noise stimulation as a means to influence motor control by “boosting” proprioception during locomotor adaptations to forces delivered by a robot. The forces increase step length unilaterally and trigger an adaptive response that restores the original symmetry. Healthy participants performed two adaptation experiments, one with stimulation applied to the hamstring muscles and one without. We found that participants adapted faster but to a lesser extent when undergoing subsensory stimulation. We argue that this behaviour is due to the dual effect that the stimulation has on the afferents encoding position and velocity that are present in the muscle spindles.

## Introduction

Subsensory mechanical and electrical noise stimulation targeting sensory receptors can alter the kinesthetic sense and lead to improved motor performance (*1–4*). This effect is understood to be related to the Stochastic Resonance phenomenon, for which small amounts of noise can improve the detection of weak signals in nonlinear threshold-based systems (*5, 6*). The noise, if correctly calibrated, increases the sensitivity of the threshold-based system, allowing for the detection of otherwise undetectable signals. An optimal level of noise maximizes this effect. Stochastic Resonance has been observed to occur in response to noise stimulation in threshold-based biological systems (*5*), in neurons (*7*) and in human sensory receptors (*8*). It is believed that noise applied mechanically or electrically to sensory receptors boosts proprioception by depolarizing the associated afferents and facilitating their firing and the transmission of the sensory information to the Central Nervous System (CNS).

Mechanical and electrical stimulation targeting joints, muscles and the soles of the feet, has been consistently shown to improve posturographic parameters during different balance exercises (*1, 2, 17, 9–16*). However, the application of this technique to lower extremity tasks different than balance has been more limited (*4, 18–21*), making it unclear whether subsensory stimulation can be successfully used also in other contexts and whether it can be selectively used to elicit desired changes in the motor behaviours associated with the performance of dynamic tasks.

Proprioceptive feedback plays a crucial role during locomotion, regulating the magnitude and duration of the activity of the various flexor and extensor muscles (*22–25*). Proprioception is also central during lower limb motor adaptations, by generating the kinesthetic component of the sensory prediction error that drives adaptation (*26*). It has been also observed that locomotor adaptation during split-belt treadmill experiments is associated with a recalibration of the perception of limb speed (*27, 28*), a process that has been attributed to the cerebellum (*29*). Given its dependence on proprioceptive input, locomotor adaptation appears to be an appropriate task to test whether the use of subsensory noise stimulation to boost proprioception affects dynamic motor control.

Based on this rationale, we tested the use of subsensory electrical current applied directly on the surface of the hamstring muscles for the purpose of altering the response of healthy individuals to a unilateral haptic perturbation delivered using a commercial robot for gait rehabilitation. The perturbation was active during the swing phase of the gait cycle and had the effect of increasing the step length of the participants’ right foot and triggering an adaptive process aiming at restoring the original step length, following a known paradigm (*30, 31*). We hypothesized that the subsensory electrical stimulation would improve adaptation performance, expressed as a faster adaptation process, by maximizing the amount of proprioceptive information available from a group of muscles involved in the adaptation process (*31*). As expected from our hypothesis, we found that the stimulation leads to faster adaptations. However, to our own surprise, we observed that the faster adaptation converged towards a longer residual step length, indicating a less complete adaptation process. The faster adaptation could be due to the increase in available proprioceptive information, as per our hypothesis, and/or from the larger target residual step length. Given this observation, these results cannot fully confirm our initial hypothesis. We theorize that our results may depend on the differential effect that the stimulation has on the different sensory receptors present in the muscle spindles, although future studies are necessary to confirm this. These results may have implications on the use of subsensory stimulation during robot-assisted gait therapy of individuals presenting with proprioceptive deficits.

## Results

### Experimental Setup, testing protocol and analyses

In this study, 30 healthy individuals (15 women, average age = 27.4 ± 3.8, average BMI = 22.9 ± 2) performed two locomotor adaptation experiments using the Lokomat device (Hocoma, Switzerland), a commercially-available robot designed for administering Robot-Assisted Gait Training (*32*). Participants performed one locomotor adaptation experiment while experiencing subsensory electrical stimulation during the whole experiment (STIM condition) and one while experiencing the stimulation only in the first 30 seconds of the experiment (NOSTIM condition). The two experiments were carried out in different days and in a random order, with the participants unaware of the condition associated with each experiment. During each locomotor adaptation experiment, participants experienced 80 steps of unperturbed walking (Baseline phase) during which the robot was fully back-driven and compensating for the interaction forces between itself and the user, followed by 80 steps of perturbed walking (Force-field phase), and finally 80 steps of unperturbed walking (After-effect phase) during which the robot was once again fully back-driven. The perturbation, whose effects have been examined in previous studies (*30, 31*), was designed as a velocity-dependent force-field acting on the right leg during the swing phase. The net effect of the perturbation was to increase the step length of the right foot (Figure 1, A). The magnitude of the force-field was dependent on the weight of the participant and was tuned so as to induce a step length increase of approximately 30% with respect to the baseline step length (*30*). During both locomotor adaptation experiments, a pair of electrodes for electrical stimulation were placed on the participants’ hamstrings (Figure 1, B). In both experimental conditions, participants received subsensory electrical stimulation consisting of zero mean Gaussian white noise with RMS amplitude value set at 90% of the sensory threshold (estimated as 191.3 ± 144.8 μA through all the experiments of all the participants). The frequency of applied stimulation was near 0 Hz to 1 kHz and was the same for all subjects. Each experiment lasted for about 10 minutes. As in previous studies based on the same paradigm (*30, 31*), the adaptation to the perturbation was quantified as the change in step length observed during the different phases of the experiments. Step length was calculated from the joint angles measured by the robot (see Materials and Methods) in a Cartesian coordinate system with origin at the hip joint of the robot. The estimated length of each step taken by study participants was normalized and expressed as percentage change with respect to the average step length observed during the baseline phase of each experiment. Adaptation and deadaptation are usually characterized by a marked exponential-like trend (*30*). For this reason, adaptation and de-adaptation behaviours were modeled by fitting exponential functions (Figure 1C and Equation 5) to the data recorded during the Force-field and After-effect phases of the experiments. The timing of adaptation and de-adaptation was estimated from the time constant of the fitted exponentials, as a mean to verify our hypothesis. We also analyzed the values of normalized step length in specific sections of the experiments. In our analyses, we first compared the changes in step length and gait biomechanics between the STIM and NOSTIM conditions, regardless of which condition was experienced first. We also analyzed differences between the two stimulation conditions by stratifying the participants in two groups, based on the order of the two experiments (STIM first or NOSTIM first).

**Fig. 1.**
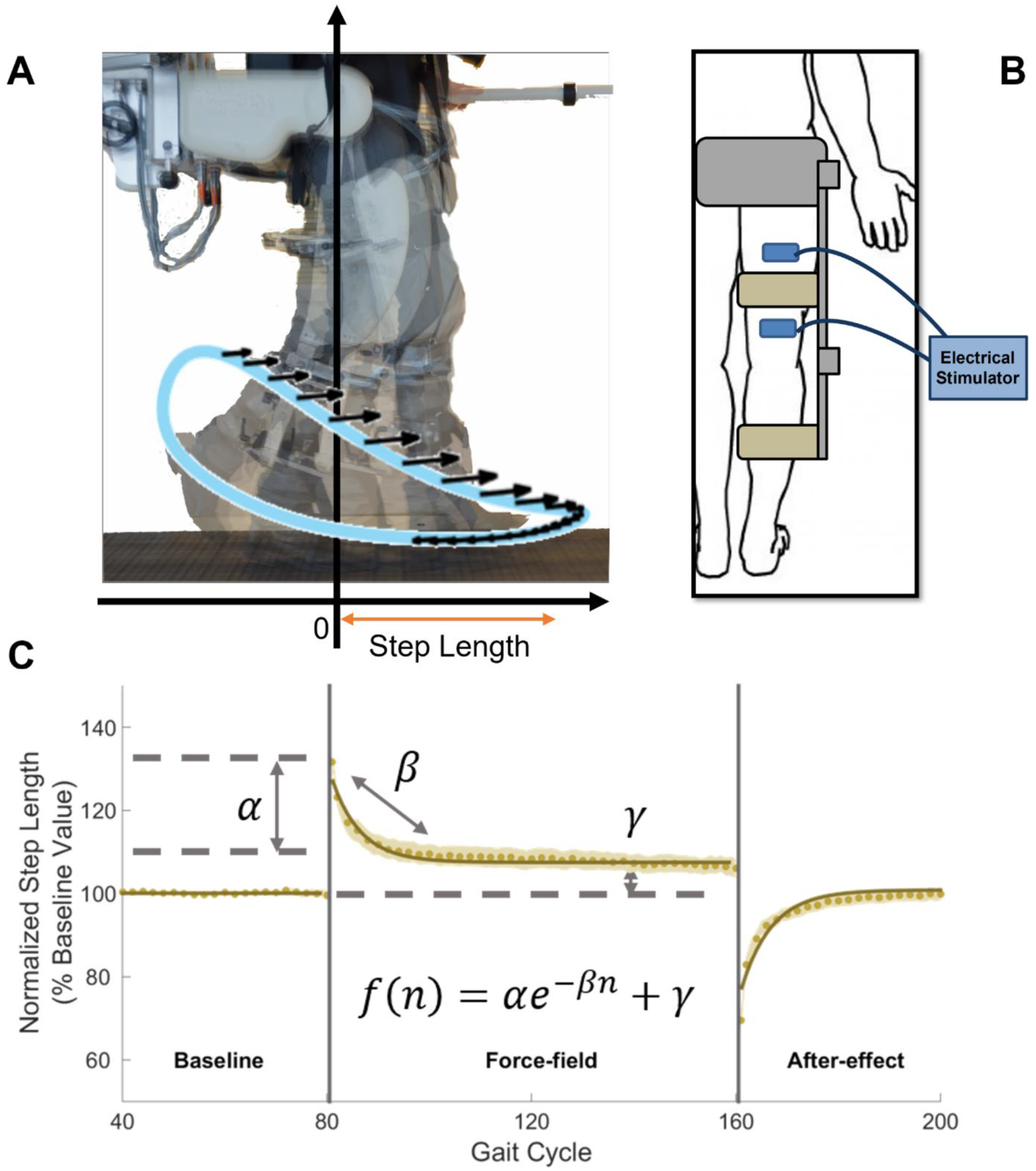
Experimental Setup and Metrics. (**A**) Lokomat setup for the motor adaptation experiment. Calculation of step length was performed in the reference system of the Lokomat, with the zero set at the mechanical center of the hip joint. The perturbing force-field (black arrows) was active during the swing phase of the gait cycle. (**B**) The electrodes for SR stimulation were placed approximately 5 cm above and below the tight upper cuff on the back of the right leg of each participant, targeting the hamstrings. (**C**) Amount and rate of adaptation and de-adaptation were calculated from the normalized step length and were modeled using the exponential function in the equation.

### Analysis of the effect of the stimulation conditions on step length adaptation

When comparing the two stimulation conditions regardless of order, we found two distinct behaviors in the adaptation plots for the STIM and NOSTIM conditions (Figure 2A). Specifically, in the Force-field phase, we observed exponential changes in step length that converged faster and towards a higher target step length in the STIM condition compared to the NOSTIM condition. This observation suggests that participants, during the STIM condition, adapted faster but to a less extent compared to the NOSTIM condition. Figure 2B shows the time constants of adaptation associated with the exponential fittings performed on the average step length data for the complete STIM and NOSTIM datasets. The number of steps needed for adaptation were estimated as 3 times the time constant *β* of the exponential function fitted to the data. From the analysis of the time constant derived from the fitted exponentials, expressed as the estimated value of steps ± 95% confidence interval, we found that participants took approximately 11.1 ± 1.6 steps (R^2^ of fitting = 0.94, rmse 0.06) to adapt to the perturbation in the STIM condition, whereas 20.1 ± 3.0 steps were needed in the NOSTIM condition (R^2^ of fitting = 0.92, rmse = 0.10). Similarly, in the After-effect phase, participants took approximately 12.6 ± 1.5 steps (R^2^ of fitting = 0.94, rmse = 0.10) to de-adapt the modified gait pattern in the STIM condition, whereas 21.1 ± 4.2 steps were needed in the NOSTIM experiment to achieve the same result (R^2^ of fitting = 0.89, rmse = 0.02). Figure 2C shows the results of the statistical analysis performed to compare the step length values observed at different points of the experiments in the STIM and NOSTIM conditions. At the first step of the Force-field phase, participants showed a normalized step length value of 132.9 ± 14.2% for the STIM condition and of 130.4 ± 10.4% for the NOSTIM condition. This difference was not significant (p = 0.185, Wilcoxon’s signed rank test with Bonferroni-Holm correction). The average step length across the last 5 steps of adaptation was 109.4 ± 12.1% for the STIM condition and 103.9 ± 9.1% for the NOSTIM condition. This difference was shown to be statistically significant (p = 0.024, Wilcoxon signed rank test with Bonferroni-Holm correction), supporting the preliminary observation that participants adapted to a lesser extent during the STIM condition. At the beginning of the After-effect phase we observed a step length of 72.5 ± 11.7% for the STIM condition and 67.6 ± 12.8% for the NOSTIM condition. This difference was statistically significant (p = 0.04, Wilcoxon’s signed rank test with Bonferroni-Holm correction) and may be explained by the smaller amount of adaptation observed for the STIM condition at the end of the Force-field phase. Figure 2D presents the analysis of the inter-subject and intra-subject variability in step length at the end of the Forcefield phase. We calculated inter-subject variability in step length as the standard deviation across participants of the average step length observed in the last 30 steps of the Force-field phase. For the STIM condition, we observed a value of inter-subject variability equal to 12.7 ± 0.9% compared to 10.3 ± 1.0% for the NOSTIM condition. This difference was shown to be statistically significant with a p < 0.001 (Wilcoxon’s signed rank test with Bonferroni-Holm correction). Intra-subject variability was calculated as the standard deviation of step length across steps observed in the last 30 steps of the Force-field phase. For this parameter, we did not observe significant differences between the STIM and NOSTIM conditions, with values of 3.4 ± 2.7% for the former and 3.4 ± 2.3% for the latter (p = 0.688, Wilcoxon’s signed rank test with Bonferroni-Holm correction). These results suggest an increase in variability between the different participants when exposed to stimulation that does not appear to be caused by an increase in the intrinsic step-by-step variability of each participant.

**Fig. 2.**
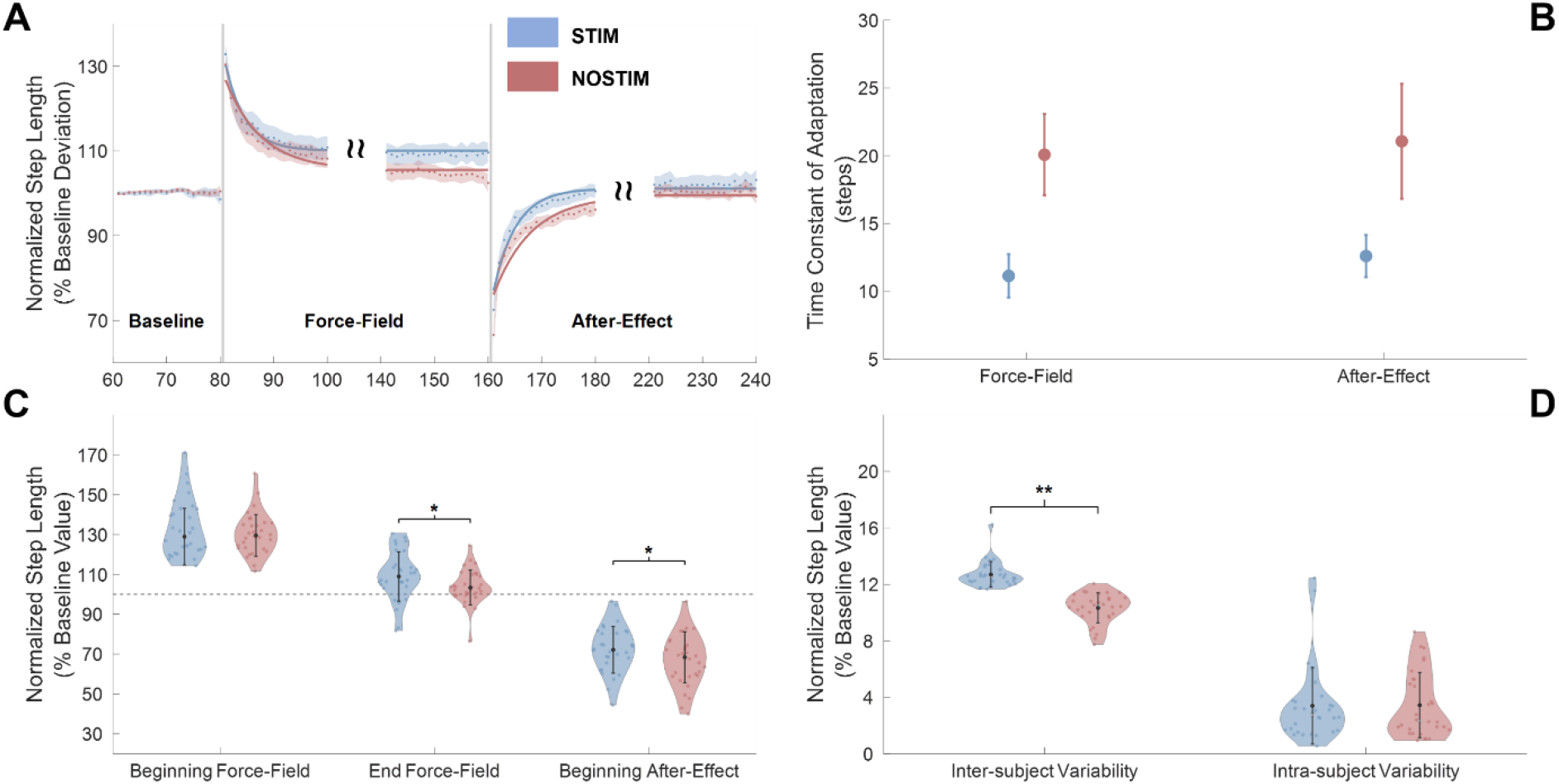
Analysis of step length adaptation, STIM vs NOSTIM conditions. (**A**) Step length changes over the course of the experiment for the STIM (light blue) and NOTISM (light red) conditions. Dots and shaded areas represent average and standard error of the normalized step length across all participants for each condition. Bold lines represent the exponential fitting calculated from the average data of each condition. Two different behaviors are clearly identifiable between the STIM and NOTISM conditions. (**B**) Time constants (in steps ± 95% confidence interval) of the exponentials estimated during the Force-field and After-Effect phases for the STIM and NOSTIM conditions. The STIM condition presents faster adaptation and deadaptation time constants. (**C**) Across-participants average ± standard deviation of step length at the beginning and end of the Force-Field and at the beginning of the After-Effect. The STIM condition presented significant higher residual step length at the end of the Force-Field and significantly lower initial After-Effect with respect to the NOSTIM condition, indicating a smaller extent of adaptation. Error bars represent median ± standard deviation across participants. (**D**) Inter-subject and intra-subject variability in step length. STIM condition is associated with a significant higher inter-subject variability, while intra-subject variability is unaltered between the two conditions. Error bars represent median ± standard deviation across participants.

### Session-dependent Analysis

Locomotor adaptations can present savings, which is a phenomenon where an adaptation experiment experienced after an earlier one is characterized by a smaller initial error and a faster adaptation speed (*33*). To account for the possibility that this phenomenon affects the results, we analyzed the dataset by stratifying participants depending on the order of the experiments. In this analysis, we split the datasets depending on whether the participants experienced the STIM or NOSTIM experiment in the first experiment and assessed each group (STIM-First and NOSTIM- First) of 15 participants separately. The results are presented in Figure 3, where blue and red indicate the STIM and NOSTIM conditions and light and dark color represent, respectively, the order of the experiment (first or second). The adaptation plots (Figure 3A) for the session-dependent analysis present adaptation trends consistent with those observed in the complete analysis. Specifically, the two STIM step length plots are “above” the respective NOSTIM datasets in the Force-field phase indicating a faster but less extent of adaptation. The analysis of the time constants confirmed this observation. The STIM-First group, in fact, presented a time constant of adaptation equal to 8.8 ± 1.9 steps for the STIM condition and 25.0 ± 3.0 steps for the NOSTIM condition (R^2^ of fittings = 0.89 for both estimations, rmse = 0.08 and 0.15 respectively). The NOSTIM-First group presented the same directional trend of STIM adaptation being faster than the NOSTIM one, with a time constant of 15.8 ± 1.8 steps for the STIM condition and of 16.9 ± 0.8 steps for the NOSTIM condition, although in this case the confidence intervals of the estimation overlap considerably (R^2^ of fittings = 0.97 and rmse = 0.03 for STIM and 0.99 and rmse = 0.05 for NOSTIM). We also observed the same behavior in the time constants of the After-effect with faster time constant associated with the STIM conditions for both groups (16.6 ± 2.6 steps, R^2^ = 0.91 for the STIM-First group, 8.8 ± 1.0 steps, R^2^ = 0.96 for the NOSTIM-First group) compared to the NOSTIM conditions (26.7 ± 4.8 steps, R^2^ = 0.89 for the STIM-First group, 19.6 ± 1.7 steps, R^2^ = 0.99 for the NOSTIM- First group). The analysis of step length showed no substantial differences across the different groups in the initial response to the perturbation. The STIM-First group presented an initial deviation of 136.1 ± 4.2% in the STIM condition and of 132.1 ± 2.7% in the NOSTIM condition (p = 0.303, Wilcoxon’s signed rank test with Bonferroni-Holm correction). Similarly, the NOSTIM- First group presented an initial deviation of 129.6 ± 2.6% in the STIM condition and of 128.7 ± 2.5% in the NOSTIM condition (p = 0.978, Wilcoxon’s signed rank test with Bonferroni-Holm correction). At the end of the Force-field phase, participants in the STIM-First group presented a residual step length of 110.0 ± 3.6% in the STIM condition and of 102.7 ± 2.5% in the NOSTIM condition. This difference, however, was not statistically significant (p = 0.0520, Wilcoxon’s signed rank test with Bonferroni-Holm correction). In the NOSTIM-First group, we observed a residual step length of 108.9 ± 2.9% in the STIM condition and of 105.8 ± 2.2% in the NOSTIM condition, although in this case the difference was not significant (p = 0.909, Wilcoxon’s signed rank test with Bonferroni-Holm correction). At the beginning of the After-effect phase, we observed, for the STIM-First group, a step length equal to 70.6 ± 3.4% in the STIM condition and equal to 62.2 ± 2.9% in the NOSTIM condition. In the NOSTIM-first condition, we observed values for step length equal to 74.4 ± 2.4% and 71.0 ± 3.2% for the STIM and NOSTIM conditions. Only the differences observed for the STIM-first group were shown to be statistically significant (p = 0.039 for the STIM-First group, p = 0.978 for the NOSTIM-First group, Wilcoxon’s signed rank test with Bonferroni-Holm correction). Analysis of inter-subject variability (Figure 3D) showed statistically significant differences between the two stimulation conditions in the STIM-First group (15.2 ± 0.9% for STIM, 8.1 ± 1.6% for NOSTIM, p < 0.001, Wilcoxon’s signed rank test) but not in the NOSTIM-First group (9.9 ± 2.0% for STIM, 11.9 ± 2.2% for NOSTIM, p = 0.454, Wilcoxon’s signed rank test). Intra-subject variability did not present statistically significant differences between conditions for both groups, with values of 3.0 ± 2.0% for STIM and 2.67 ± 1.6% for NOSTIM in the STIM-First group (p = 0.978, Wilcoxon’s signed rank test) and 3.0 ± 4.1% for STIM and 3.0 ± 2.0 for NOSTIM in the NOSTIM-First group (p = 0.909, Wilcoxon’s signed rank test).

**Fig. 3.**
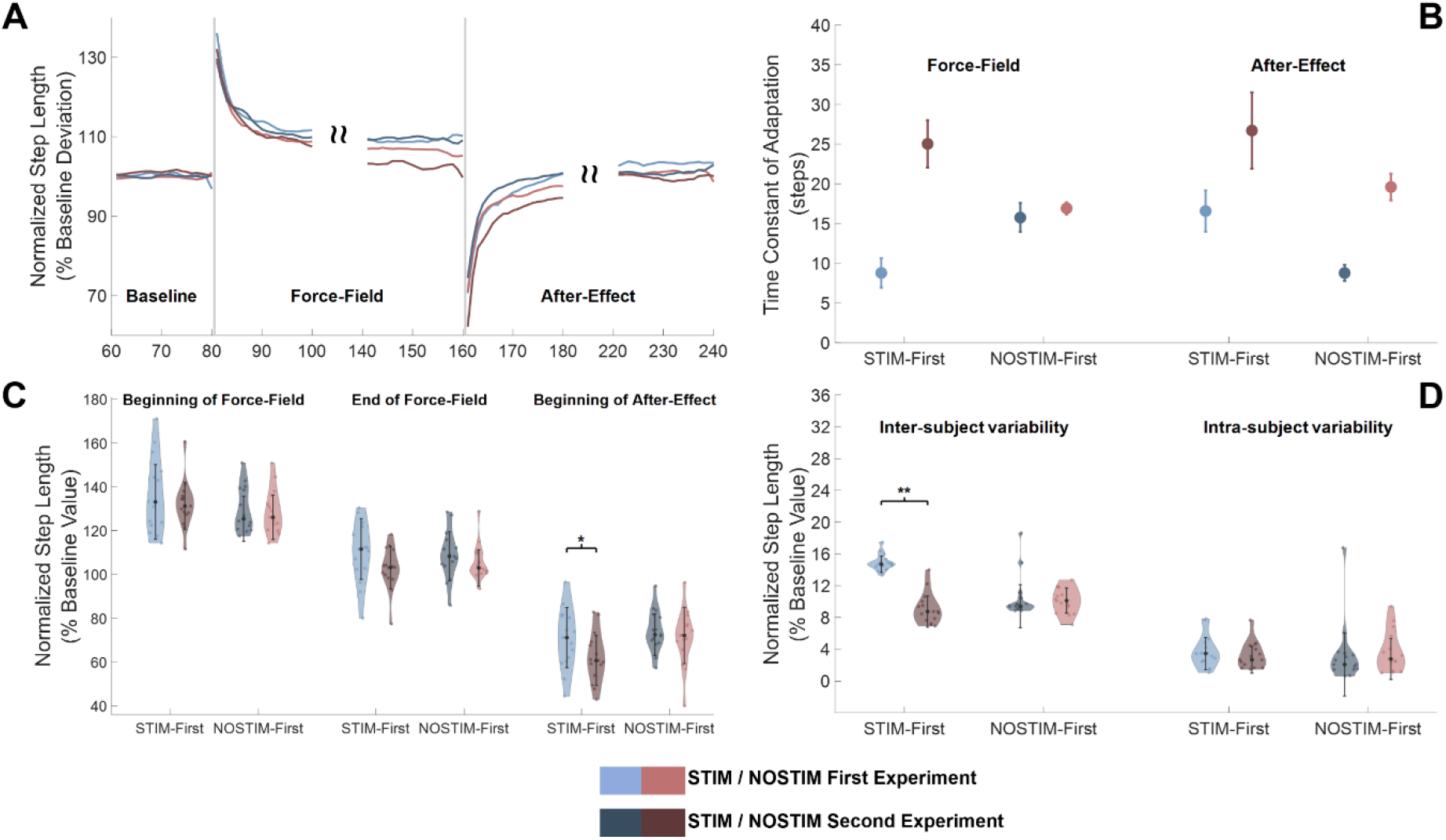
Session-dependent analysis of step length adaptation, STIM-First and NOSTIM-First groups. (**A**) Adaptation plots for the STIM (blue) and NOSTIM (red) conditions depending on the order of the experiment. Lighter color indicates that a condition has been experienced in the first experiment, while darker color in the second. Dots and shaded areas represent average and standard error of the normalized step length across all participants for each session/condition. Both STIM conditions present faster and less extended adaptation with respect to the respective NOSTIM conditions. (**B**) Time constants (in steps ± 95% confidence interval) of the exponentials estimated during the Force-field and After-Effect phases for the STIM and NOSTIM conditions of the STIM-First and NOSTIM -First groups. The STIM conditions present faster adaptation and de-adaptation time constants in both groups. (**C**) Across-participants average ± standard deviation of step length at the beginning and end of the Force-Field and at the beginning of the After-Effect. The STIM conditions presented higher residual step length at the end of the Force-Field and significantly lower initial After-Effect with respect to the respective NOSTIM conditions. This result was statistically significant (p<0.05, Wilcoxon signed rank test) only for the STIM-First group. Error bars represent median ± standard deviation across participants. (**D**) Inter-subject and intra-subject variability in step length. STIM condition for the STIM-First group presented a statistically significant higher inter-subject variability. Error bars represent median ± standard deviation across participants.

### Effect of the Stochastic Resonance stimulation on the joint kinematics and kinetics at the hip and knee during adaptation

We analyzed the joint biomechanics to identify a possible cause of the differences in step length adaptation between the STIM and NOSTIM conditions. The average joint angles at the end of the adaptation period are similar in both conditions. The higher residual step length observed in the STIM condition compared to the NOSTIM condition is mainly explained by a slight anticipation in the hip and knee angle patterns observed from about 40% of the gait cycle on. In particular, we observed an anticipation of the position of the peak hip flexion during swing in STIM with respect to NOSTIM (Table S1 in Supplementary Materials), which however was not significant after adjusting for multiple comparisons. We did not observe substantial differences in the measured torques at both the hip and knee between baseline and the last steps of adaptation in both the STIM and NOSTIM conditions. It needs to be pointed out that the measured torques are the interaction torques between the user and the robot measured by the Lokomat sensors and do not capture completely the torques generated at the hip and knee during gait. We observed small, non-significant, differences between the STIM and NOSTIM conditions for the hip torque, mostly characterized by a higher negative peak during late stance in STIM and an anticipated torque profile, which agrees with what observed in the kinematics.

At the knee, for both the STIM and NOSTIM conditions, we observed an increased peak knee torque during swing, which was higher for STIM compared to NOSTIM. In STIM, we also observed a lower negative peak during terminal swing, which appeared delayed in NOSTIM. Both these results were not statistically significant (table S1 in Supplementary Material). Adaptation to the perturbation is associated with an increase in joint power generated against the machine during the swing phase at both the hip and knees. At the knee level, in particular, a peak of generated power is present in late swing where normally a power absorption is observed, in a behavior that reflects the participants rejection of the step lengthening perturbation. Small differences, not statistically significant, were observed between the two conditions, with STIM associated with an anticipated hip power generation profile and a higher knee power peak during late swing. We estimated the power required to counteract the perturbation as the difference between the average power calculated during the baseline phase and that calculated in the last 5 steps of the adaptation phase, minus the power attributed to the perturbation itself (see Material and Methods for details). We observed, for both hip and knee and in both STIM and NOSTIM conditions, that participants exerted additional power at the beginning of the stance phase (loading-response) and during mid/terminal swing, when the perturbation is active. For both conditions the amount of additional power was higher at the knee with respect to the hip. At the hip we observed a higher amount of power exerted by the participants during the NOSTIM condition with respect to the STIM one, although this difference was shown to be not significant (see Supplementary Material, Figure S1). At the knee, on the other hand, participants presented a higher additional power during the STIM condition with respect to the NOSTIM condition (Figure S1 Supplementary Material), but also this difference was not found to be statistically significant.

**Fig. 5.**
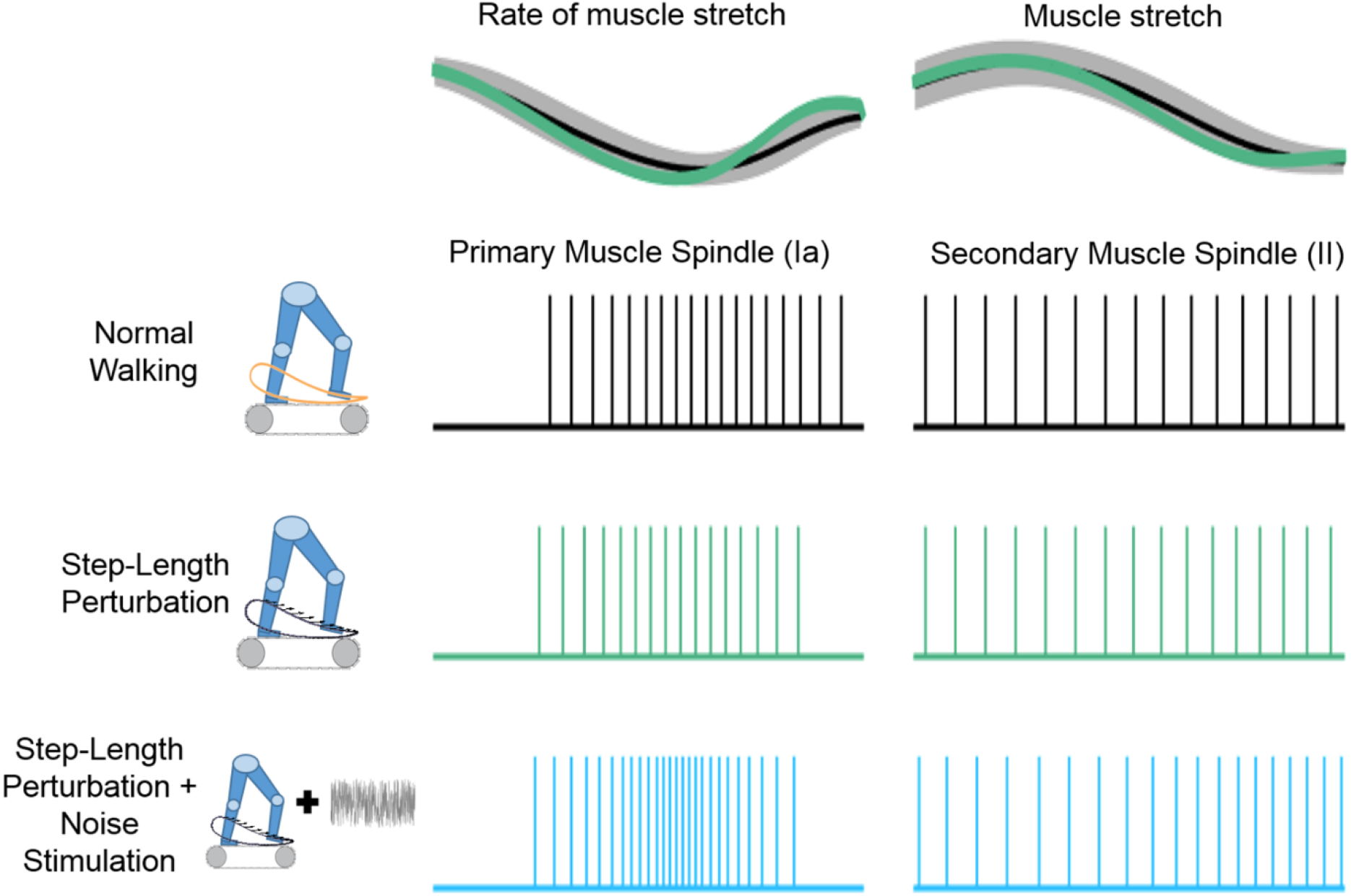
Theoretical model of the effect of subsensory electrical stimulation on the spindles of the hamstring muscles during the locomotor adaptation experiments. The response of primary muscle spindles (afferent Ia) is modeled as a firing depending on the rate of muscle stretch (modeled, for the hamstring muscle during swing, as the derivative of the knee angle, top plot, left), while the response of the secondary muscle spindles (afferent II) is modeled as a firing directly dependent on the amount of muscle stretch (modeled as the changes in knee angle, top plot, right). The perturbation (third row) leads to increased firing rate during late swing phase for both the Ia and II afferents. The effect of the subsensory stimulation is that of increasing the depolarization level of the spindles, affecting the firing/stretch relationship for both the Ia and II receptors. During subsensory electrical stimulation higher firing rates are associated to the same stretch and rate of stretchy induced by the perturbation, possibly altering the active response of the muscle through spinal and supra-spinal circuits.

## Discussion

This study investigated the effect of subsensory electrical noise stimulation applied to the hamstring muscles during a robot-based locomotor adaptation experiment. Our results revealed that subsensory stimulation directly altered the adaptation process so that in the experiments where the stimulation was applied, individuals adapted faster but to a lesser extent compared to experiments during which no stimulation was delivered. We also saw that this effect is bigger in the group that received the stimulation in their first experiment. Our hypothesis was that the subsensory electrical stimulation would increase the sensitivity of the muscle spindles and thus accelerate the adaptation process by increasing the amount of proprioceptive information available to the CNS when processing and responding to the perturbation.

Although our results seem to confirm this hypothesis the reasons behind the accelerated adaptation that we observed are not completely clear. In fact, the observed increase in adaptation speed could be directly due to the stimulation or could be secondary to the larger kinematic error observed at the end of the force-field period in the STIM condition. In the first case, the direct effect of the peripheral stimulation would be that of accelerating the adaptation process, while in the second case, the smaller number of steps required to reach the adapted plateau would be a by-product of the fact that the adaptation target is closer to the original deviation induced by the perturbation. In the following, we will discuss these two possible mechanisms. During motor adaptations to perturbations that induce dynamic changes in the walking environment, the CNS modifies the motor plan so as to anticipate the expected effects of the perturbation with the aim of minimizing the sensory prediction error associated with the kinematic error. In our experiments we induced a kinematic error in step length that leads to an asymmetric gait pattern. Adaptation in this case is likely associated with the restoration of the normal relationship between the center of mass and the boundaries of the base of support to maintain a stable gait pattern (*30, 34*). Emken and colleagues showed that motor adaptations during walking can be described as a tradeoff between the kinematic error caused by the perturbation and the effort required to reject it (*35*). The same authors demonstrated that the rate of adaptation, herein measured by the number of steps needed for the adaptation process to converge, depends on the magnitude of the perturbing force-field (*36*), where stronger perturbations cause bigger kinematic errors and thus faster adaptation processes.

Rate of adaptation has also been shown to depend on the presence and structure of previous exposures to the perturbation (*33*). Specifically, Malone and colleagues showed that individuals who had experienced multiple exposures to the same perturbation in the same day presented faster adaptations in the exposures following the first one, in a process often referred to as “savings” (*33*). We designed our experiments so as to limit the possible effects of savings by performing the adaptation experiments over non-consecutive days. Savings, however, have been observed also in case of experiments performed months apart (*37*).

At a first glance, our data surprisingly showed a pattern opposite to the one observed by Malone. Our data shows that locomotor adaptation experiments performed in the second session (across both stimulation conditions) presented longer adaptation time constants with respect to those performed in the first session (Figure S2 in Supplementary Material). However, this behavior appears to be driven by the group who received the stimulation in their first experimental session. Although we observed consistently faster but less “complete” adaptation in the STIM compared to the NOSTIM condition regardless of the order of the experiment, the STIM-first group presented the fastest learning time constant among session/conditions during STIM and the slowest during NOSTIM. This result may suggest that the stimulation is more effective if the task has not been experienced recently and that having first adapted with a “boosted” proprioceptive input disrupts a subsequent adaptation, possibly by disrupting the sensory recalibration process associated with locomotor adaptation (*27*). This interpretation, although plausible, needs to be confirmed through specifically designed experiments.

The main proxies of the differences we observed between the step length adaptation processes in the STIM and NOSTIM conditions are the differences we observed in the kinematics and kinetics of the participants at the hip and knee joints. In the STIM condition, participants showed trends of slightly anticipated hip and knee angle patterns and increased knee torque and power generated during the late swing phase compared to the NOSTIM condition (Figure 4). These results show a difference in power exerted at the knee between the STIM and NOSTIM conditions (Figure 4 and S1). The behaviour observed in the STIM condition is in accordance with an anticipation in negative knee torque production in swing (when the perturbation is active) and thus consistent with a higher (and possibly anticipated) activity of the hamstring muscles that are the main drivers of the negative contribution to the active knee torque in terminal swing, and that are the muscles we observed to contribute to the rejection of this particular perturbation (*31*). While the results of our experiments cannot directly confirm that the stimulation ultimately leads to an increased activity of the hamstrings, since EMG data was not recorded, this plausible result, if verified, would clearly indicate that subsensory stimulation can be used to selectively alter the activity of different muscles during the performance of dynamic tasks.

**Fig. 4.**
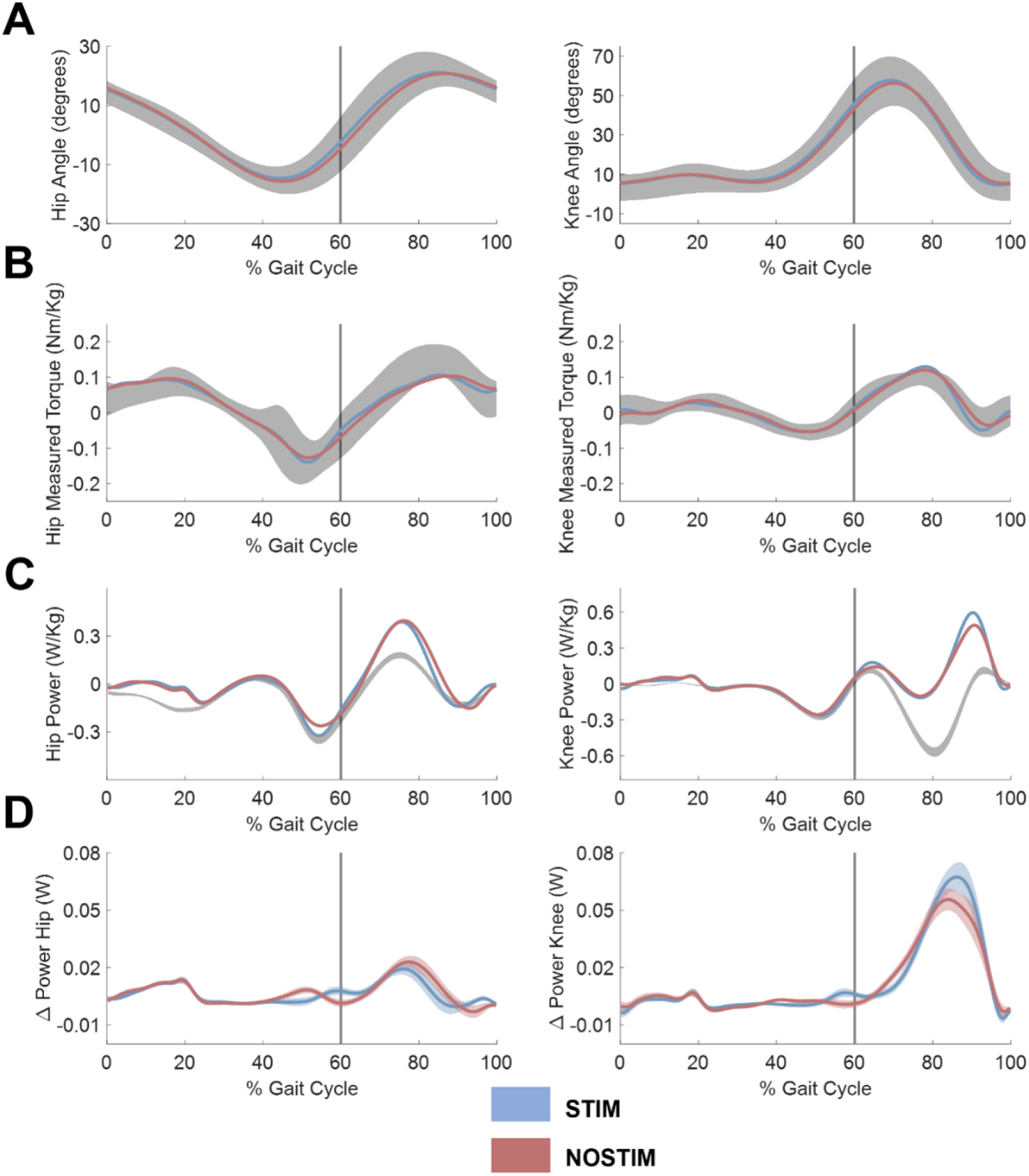
Hip/knee kinematics and kinetics during baseline and late adaptation. (**A**) Across-participants average hip (left) and knee (right) angles during baseline (shaded area represents the average ± standard deviation across all participants) and at the end of adaptation for the STIM (blue line) and NOSTIM (red line) groups (each line represents the average across participants). (**B**) Across-participants average hip (left) and knee (right) measured torques during baseline (shaded area represents the average ± standard deviation across all participants) and at the end of adaptation for the STIM (blue line) and NOSTIM (red line) groups (each line represents the average across participants). The torques are measured as the interaction torques between the individuals and the Lokomat exoskeleton and do not correspond to the joint torques. (**C**) Across- participants average hip (left) and knee (right) power during baseline (shaded area represents the average ± standard deviation across all participants) and at the end of adaptation for the STIM (blue line) and NOSTIM (red line) groups (each line represents the average across participants). The power is calculated from the measured torques and do not correspond to the actual joint power. (**D**) Average ± standard deviation of the additional power (calculated as the difference between the power at the end of adaptation and the power at baseline after removing the power associated to the pertubration) for the STIM (blue) and NOSTIM (red) conditions at the hip (left) and knee (right). The STIM condition is characterized by a slightly lower peak hip additional power, and by a significantly (see Figure S1) higher additional power at the knee level. In all eight plots the vertical grey line represents the estimated point of transition between stance and swing phases of the gait cycle (toe-off).

To get a better understanding of the processes potentially leading to our results, it is perhaps useful to consider the theoretical effects that the subsensory electrical stimulation has on the muscle spindles during movement (Figure 5). Muscle spindles are intrafusal fibers parallel to the skeletal muscle fibers that encode information regarding the rate of muscle stretch (group Ia afferents) and the amount of muscle stretch (group II afferents) during active and passive contractions (*38*). Specifically, the firing rate of group Ia afferents increases with the rate of muscle stretch, while that of group II afferents increases with the amount of stretch. Considering the hamstring muscles during the mid-to-terminal swing phase of the gait cycle, where the muscle is lengthening due to knee extension while contracting eccentrically, we can hypothesize the dynamic behaviour of the stretch to be roughly proportional to the dynamical behaviour of the knee angle. This gross approximation, that omits for simplicity these muscles’ length dependency on the hip angle, which, however, does not change substantially in late swing, is herein used to emphasize the role that the hamstring muscles have in terminal swing, where they contribute to the deceleration of knee extension.

A perturbation like the one we applied in our experiment directly modifies the dynamic stretching of the hamstring muscles. As the perturbation increases step length mostly by exerting a knee-extending torque during swing, it induces a faster and higher stretch in the hamstring muscles before its compensation, thus driving higher firing rates in both the Ia and II receptors (Figure 5), with respect to their firing patterns during baseline walking. Small electrical currents acting on the spindles theoretically lead to a partial depolarization of the receptors membrane (*2, 8*), bringing it closer to its firing threshold. This effect should facilitate the firing of the receptors and possibly translate in both the Ia and II receptors firing at a higher firing rate given the same rate and amount of stretch compared to their normal firing patterns. Thus, in this theorization, the depolarization disrupts the natural coding between the firing rate of the Ia and II receptors and the rate/amount of muscle stretch in the hamstrings, increasing the firing rate during the portion of the gait pattern where the muscle is lengthening.

This theoretical framework suggests a coupled effect of the stimulation on the rate of adaptation. The increased sensitivity of Ia receptors could, in fact, alter the perceived magnitude of the perturbation, leading the CNS to predict a higher biomechanical error, earlier during the swing phase. This effect could drive an anticipated higher activity in the hamstring muscles while compensating for the effects of the perturbation that would cause a faster adaptation similarly to what has been observed by Emken and colleagues (*36*). At the same time, the disruption of the normal firing/stretch relationship in the II receptors could alter the perceived level of stretch thus modifying the perceived target adapted muscle length, inhibiting the activity of the hamstring earlier than in the non-stimulated condition. In the context of this interpretation, the overall faster rate of adaptation observed in the STIM condition would be both a primary effect of the stimulation and a secondary effect of the larger kinematic error observed at the end of adaptation. Our results support but do not confirm this theoretical framework, that needs to be verified through specifically designed experiments involving the recording of both spindle activity and surface EMG of the hamstring.

It is worth noticing that the STIM condition is also associated with a higher inter-subject step length variability that is not dependent on an increase in intra-subject variability. This behaviour suggests a participant-specific response to the stimulation. For each participant, we selected the RMS value of the stimulating noise current equal to 90% of the sensory threshold. This value, selected based on the response of the skin nociceptors, may lead to different degrees of deeper muscle spindle depolarization across participants (also considering the differences, across participants in leg size, muscle tone and fat distribution), thus it may be “more optimal” for some individuals and less for others. This interpretation about the cause of the observed variability, could suggest the presence of a Stochastic Resonance effect (*5*)(*6*) associated with the noise stimulation, as suggested by several previous studies in literature (*1–4, 8, 10, 18*), where an optimal level of noise maximizes the operations of a threshold-base system, while smaller and higher levels of noise yield worse results. Future experiments could explore methods to optimize the stimulation level for an individual based on muscle spindle sensitization rather than skin sensation.

These results show that subsensory electrical stimulation directly modifies motor control during the performance of dynamic tasks, and that the observed kinematic effects are theoretically supported by a modification in activity of the stimulated muscle. Our findings could have interesting applications in gait rehabilitation. Our results, in fact, suggest that subsensory electrical stimulation can be used to modulate the activation of the targeted muscles in response to a stretch. Although it is not easy to predict if individuals with impaired proprioception would respond to the stimulation in the same way as we observed in healthy individuals, the ability to selectively modulate muscular response during training through peripheral proprioceptive enhancement could theoretically be useful in promoting targeted and individual-specific changes in muscular activity in response to the forces that the robot exerts on the leg.

## Materials and Methods

### Participants and ethics statement

30 healthy individuals (15 women, average age = 27.4 ± 3.8, average BMI = 22.9 ± 2) participated in this study. Among the participants, 6 had experienced the Locomotor adaptation experiment at least once before the beginning of the study, but each more than one month before the first experimental session. All procedures were performed at the Spaulding Rehabilitation Hospital in Boston. All experimental protocols were approved by the Spaulding Rehabilitation Hospital Internal Review Board.

### Experimental Setup and Sessions

The study consisted of three experimental sessions. All experimental sessions were performed using the Lokomat (Hocoma AG, Switzerland) exoskeleton. The Lokomat consists of a lower limb exoskeleton attached to a frame that allows for actuation of hips and knees while the participant walks on a treadmill. The Lokomat system allows for Body Weight Support, although this feature was not used during the experiments. The Path Control modality allows participants to naturally control the timing of their gait while using the machine (*39*). This is achieved by estimating, at every instant, the current phase of the gait cycle by comparing the actual joint position and velocities with a pre-determined pattern using the algorithm developed by Aoyagi and colleagues (*40*). For each participant, the experimental sessions were held on different days, with a maximum of 10 days between the first and the last session and at least one day of rest between consecutive Locomotor adaptation experiments.

The first experimental session was used to find the settings for correctly fitting the person inside the Lokomat. During the first session participants walked while attached to the Lokomat for approximately 10 minutes and were familiarized with walking while attached to the device. During the second and third sessions the participants underwent the Locomotor adaptation Experiments, once while experiencing continuous subsensory electrical stimulation during the whole experiment (STIM) and once with electrical stimulation only in the first 20 seconds of the experiment (NOSTIM). The order of the STIM and NOSTIM experiments was randomized across participants with 15 participants (STIM-First group) performing the STIM experiment in the second session (first Locomotor adaptation Experiment) and 15 participants (NOSTIM-First group) performing the STIM experiment in the third session (second Locomotor adaptation Experiment).

### Locomotor adaptation Experiment

After the initial setup, participants went through a *pre-baseline* part that consisted of two phases. In the first phase, participants were asked to walk for 90 seconds at 3 km/h with the machine back- driven (thus interaction forces between the Lokomat and the participant were minimized in order to give the impression of “free walking”). The data recorded during this period were used to calculate the Generalized Elasticities (*41*) that allow for the minimization of the device loading during operations. In the second phase participants were asked to walk for 120 seconds in back- driven mode (but with the newly updated elasticity) in order to record their baseline walking pattern. This baseline pattern was used as reference for the Aoyagi algorithm. After the *pre-baseline* part, which was repeated in each experimental session, participants performed the Locomotor adaptation experiment consisting of four consecutive blocks:

- **Catch trials:** 180 gait cycles, with the machine in back-driven mode. The perturbation was activated for 10 random single steps (catch trials) interspersed at least 8 steps between one another. These catch trials were used to estimate the “free” response to the perturbation, before adaptation can take place.
- **Baseline:** 80 gait cycles, with the machine in back-driven mode.
- **Force-Field:** 80 gait cycles, with the machine perturbing the participant’s gait pattern through a velocity-dependent force field applied to the right leg during swing phase.
- **After-Effect:** 80 gait cycles with the machine in back-driven mode again.

Each Locomotor adaptation experiment lasted for about 10 minutes. During all the phases of the experiment (pre-baseline and adaptation experiment) the treadmill of the Lokomat was set to 3 km/h and participants were asked to follow the beat of a metronome pacing them at one full gait cycle every 1.4 seconds (85.7 bpm). All experimental procedures, including the setup of the Lokomat and the perturbation (see below), were identical to the X experiment performed in our previously published study (*30*).

### The perturbation

The perturbation was designed as a velocity-dependent force field acting unilaterally on the right leg. The perturbation was active during the swing phase of the right foot and aimed at increasing the step-length of the participant experiencing it (see Figure 1). The perturbation force was generated by the robot according to the following equation:

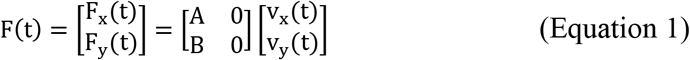

where *F_x_* and *F_y_* represent the antero-posterior (*x*) and vertical (*y*) components of the perturbing force acting on the foot, and *V_x_* and *V_y_* represent the *x*- and *y*-components of the foot velocity as reconstructed using the following Jacobian:

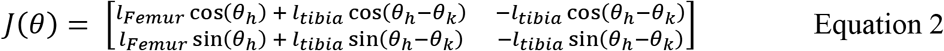

where *l_Femur_* is the length of the femur, *l_Tibia_* is the length of the tibia, and *θ* is the vector of the joint angles *θ_h_* at the hip and *θ_k_* at the knee.

The damping coefficients were calculated as

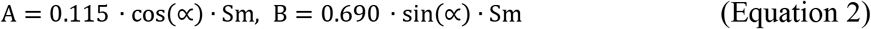

Where ∝ is the angle of the perturbation and *Sm* is the participant’s mass in kg. For this experiment the angle was set at 8°. This angle was found, in our previous experiments (*30*), as the angle at which the effect of the perturbation only act in the positive step length direction, without affecting the step height. The constant values in A and B were also empirically determined in the same experiment as values able to induce approximately a 30% increase in step length with respect to the baseline trajectory. The perturbation was applied only during the swing phase starting at the point of the gait cycle when the estimated foot velocity in the antero-posterior direction turned from negative to positive (V_x_(t) > 0| v_x(t-1)_ < 0) and ending at the point when the position of the foot in the antero-posterior direction turned from positive to negative (x(t) < 0|x(t-1)>0). The zero of the foot position in the antero-posterior direction was set at the center of the hip joint (Figure 1, Panel A).

### Data Analysis

The Lokomat system is equipped with 4 position and 4 torque sensors measuring respectively the joint angles and the interaction torques between the machine and the user at the hips and knees for both legs. The torque exerted by the participant during the experiments was calculated by subtracting the perturbation torques (calculated from Equation 4) from the interaction torques measured by the sensors. Since the effect of the perturbation administered during the Locomotor adaptation experiment was that of increasing the step length of the participants, adaptation and deadaptation were evaluated for each participant in term of changes in the normalized value of step length. Step length was calculated from Equation 3 as the maximum value observed for the *x* variable during each gait pattern. The zero step-length value was set at the center of the hip joint (Figure 1, panel A). For each experiment of each participant, the step length associated with each step was normalized by dividing it by the average step length during the Baseline part of the experiment. Locomotor adaptation is commonly characterized by an exponential change in the kinematic/kinetic metric of interest (*30, 31*). We observed a similar behavior also in the Locomotor adaptation Experiments performed in this study (Figure 1, Panel C). Exponential fitting was applied on the force-field and after-effect sessions of the experiments according to the following formula:

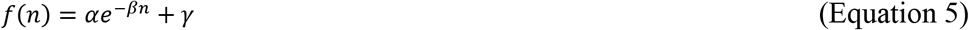

For the Force-field phase the parameter *α*, that represents the initial response to the perturbation, was estimated as the average of the deviations observed during the 9 single-step perturbations the participants experienced during the catch trial session of the experiment. The parameters *β* and *γ*, that represent respectively the time constant of the adaptation and the residual deviation at the end of the force-field phase, were estimated using a least-squares algorithm. For the after-effect phase, all the three parameters *α, β* and *γ*, were estimated, with 95% confidence intervals, using a leastsquares algorithm. Exponential fitting was applied to the normalized step length patterns averaged across all participants for both conditions (STIM and NOSTIM) and across groups of participants stratified depending on the whether they performed the STIM or NOSTIM experiments as the first Motor Adaption Experiment (STIM-First and NOSTIM-First groups). For all the analyses, the quality of the fitting was evaluated using the R^2^ parameter and the root mean squared error. The analysis of adaptation speed, estimated from the fittings as three times the value of the parameter *β* was used to verify our hypothesis that the stimulation leads to faster adaptation.

A series of additional analyses were run to characterize stimulation-dependent differences in the patterns of adaptation. Statistical analyses were performed to assess for significant (p < 0.05, after correction for multiple comparisons) differences between the step length values in different sessions of the experiments for both STIM and NOSTIM conditions. This analysis was based on Wilcoxon’s signed rank test with Bonferroni-Holm correction that has been used to compare the values of step length between the STIM and NOSTIM conditions in three specific points of the experiments. Specifically, we analyzed: a) the first step of Force-Field, to investigate the presence of differences in initial response to the perturbation between the two conditions; b) the average of the last five steps of Force-field, to assess for differences between the two conditions when the adaptation is complete; c) the first step of After-Effect, to assess for differences in the starting point of the deadaptation period.

We evaluated inter-subject and intra-subject variability in step length at the end of the force-field, when adaptation is supposed to be complete. The former measures the variability observed in each condition across the different participants, the latter measures the step-to-step variability of each participant for each condition. Inter-subject variability was calculated as the standard deviation (across participants) of the average step length calculated in the last thirty steps of the force-field phase. Intra-subject variability was instead calculated as the average standard deviation observed for each participant in the last thirty step of the force-field phase. Power was estimated, for both the hip and knee, from equation 6:

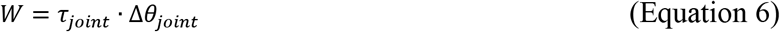

Where *τ_joint_* is the interaction torque measured at the joint and Δ*θ_joint_* is the joint angular velocity. We analyzed the differences among STIM and NOSTIM conditions in the power *W_add_* needed to counteract the perturbation. This value was estimated, for each joint, as:

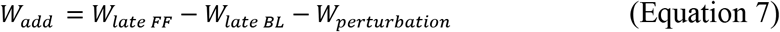

Were *W_late FF_* is the power estimated in the last ten steps of the Force-field phase, *W_late BL_* is the power estimated in the last ten steps of the Baseline phase and *W_perturbation_* is the power that is attributed to the perturbation. This value was estimated as:

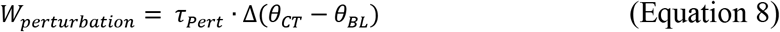

Where *τ_Pert_* is the perturbing torque calculated in Equation 4 and *θ_CT_* and *θ_BL_*represent, respectively, the joint angles during the catch trials (where adaptation to the perturbation is not yet present) and the average angle measured during the Baseline condition.

### Electrical Stimulation

During both the STIM and NOSTIM experiments electrical noise stimulation was applied through surface electrodes placed above the hamstring muscles (Figure 1, Panel B). The electrodes were rectangular self-adhesive gel pads (Axelgaard Mfg. Co., Ltd., Fallbrook, CA, USA) placed approximately 5 cm above and below the upper cuff of the Lokomat. The electrical stimulation was Gaussian White Noise with zero mean and fixed root mean square (RMS) electrical current and was generated using a custom-developed isolated stimulator commanded via a custom software application. The frequency of the stimulation was between close to 0 Hz and 1 kHz. The RMS value of the electrical noise was set for each trial of each participant at 90% the RMS corresponding with the sensory threshold of the participant, where the sensory threshold is defined as the minimum RMS for which the participants were able to feel the stimulation on their skin. The sensory threshold was estimated for each participant before the beginning of each experiment. Specifically, after the participants were secured to the Lokomat system and before the beginning of the pre-baseline walking trials, the electrodes were placed on the skin and the procedure for defining the sensory threshold was initiated. A first check on the placement of the electrode was performed by applying a noise current with an RMS of 500 μA (that represented the upper limit of the possible RMS values), to check if participant could feel the stimulation and to show them the tickling sensation associated with a noise stimulation above the sensory threshold level. Next, we set the RMS of the noise stimulation at 20 μA and increased the value by 10 μA every 10 seconds until the participant reported a tickling sensation on the skin. The RMS of the electrical noise stimulation for the trial was then set at 90% this value. During the NOSTIM condition the subsensory electrical noise stimulation was applied only at the very beginning of the experiment, during the first 20 seconds of the Catch Trials phase. In the STIM condition the stimulation was applied during the whole experiment. Participants were told, before each session, that electrical noise stimulation may be applied to them during the Lokomat experiment. Since the stimulation was always below sensory threshold, participants were effectively blinded on the experimental condition during each experimental session.

## Competing interests

JN is currently a technical advisor to Accelera, a United States based early state company looking to develop products using stochastic resonance stimulation to improve neurological deficits in patients. PB has received grant support from the American Heart Association, the Department of Defense, the Michael J Fox Foundation, the National Institutes of Health (NIH), the National Science Foundation (NSF), and the Peabody Foundation including sub-awards on NIH and NSF SBIR grants from Barrett Technology (Newton MA), BioSensics (Watertown MA) and Veristride (Salt Lake City UT). He has also received grant support from Emerge Diagnostics (Carlsbad CA), MC10 (Lexington MA), Mitsui Chemicals (Tokyo Japan), Pfizer (New York City NY), Shimmer Research (Dublin Ireland), and SynPhNe (Singapore). He has served on the Advisory Board of SwanBio (Boston MA). PB serves in an advisory uncompensated role the Michael J Fox Foundation the NIH-funded New England Pediatric Device Consortium, and the Walking Tall-PD clinical trial carried out by Neuroscience Research Australia. He also serves in an uncompensated role on the Scientific Advisory Boards of ABLE Human Motion (Barcelona, Spain), FormSense (San Diego CA, USA), Hocoma AG (Zurich, Switzerland), and Trexo (Toronto, Canada).

## Supplementary Materials for

**Fig. S1.**
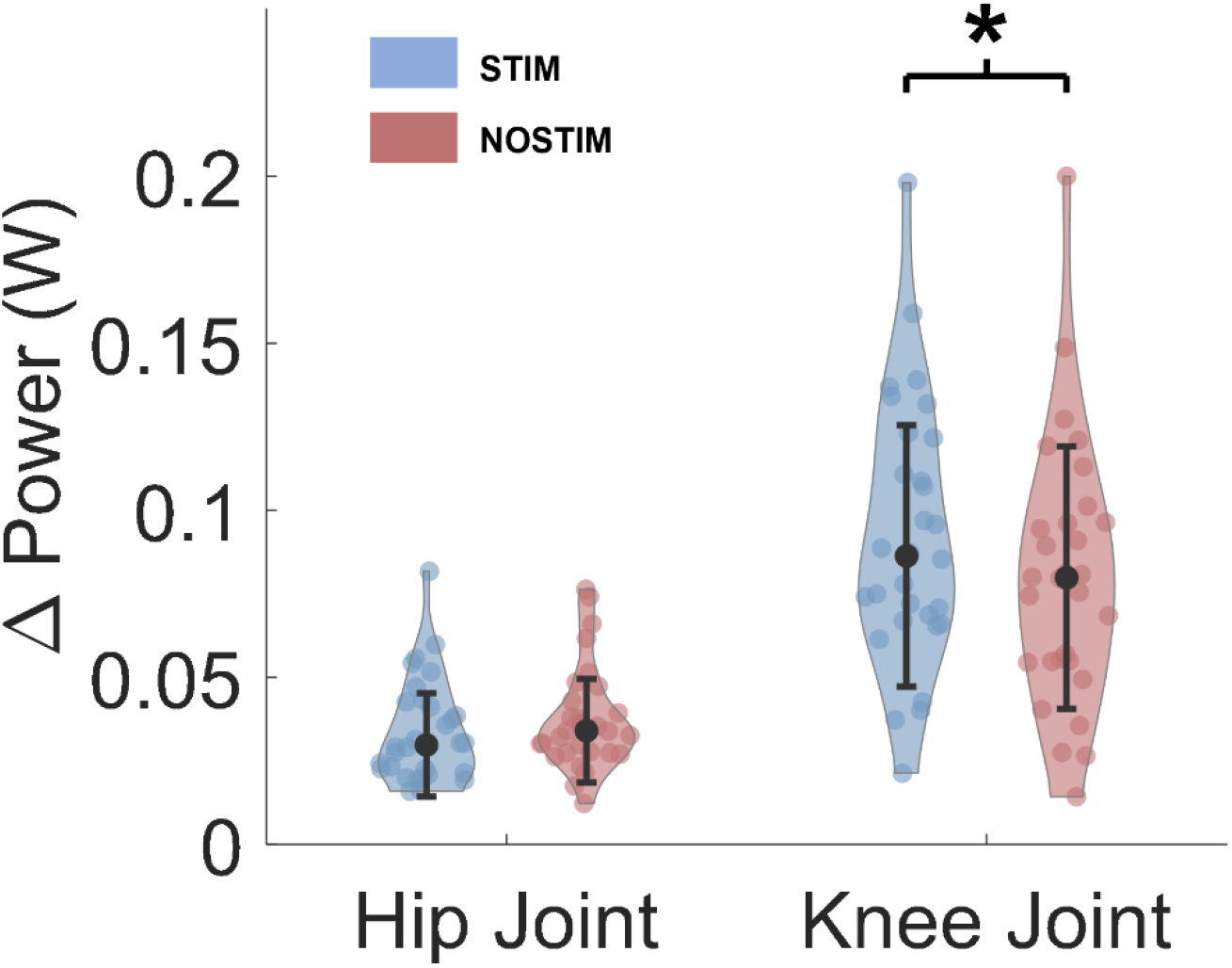
Analysis of the peak additional Power in the STIM and NOSTIM conditions. Error bars represent median ± standard deviation across participants. * indicates statistically significant differences between the two conditions with p<0.05 estimated by Wilcoxon’s signed rank test. Blue violin plots represent the STIM group, red violin plot represent the NOSTIM group. We observe significant changes in peak additional mechanical work between the two conditions only at the knee level.

**Fig. S2.**
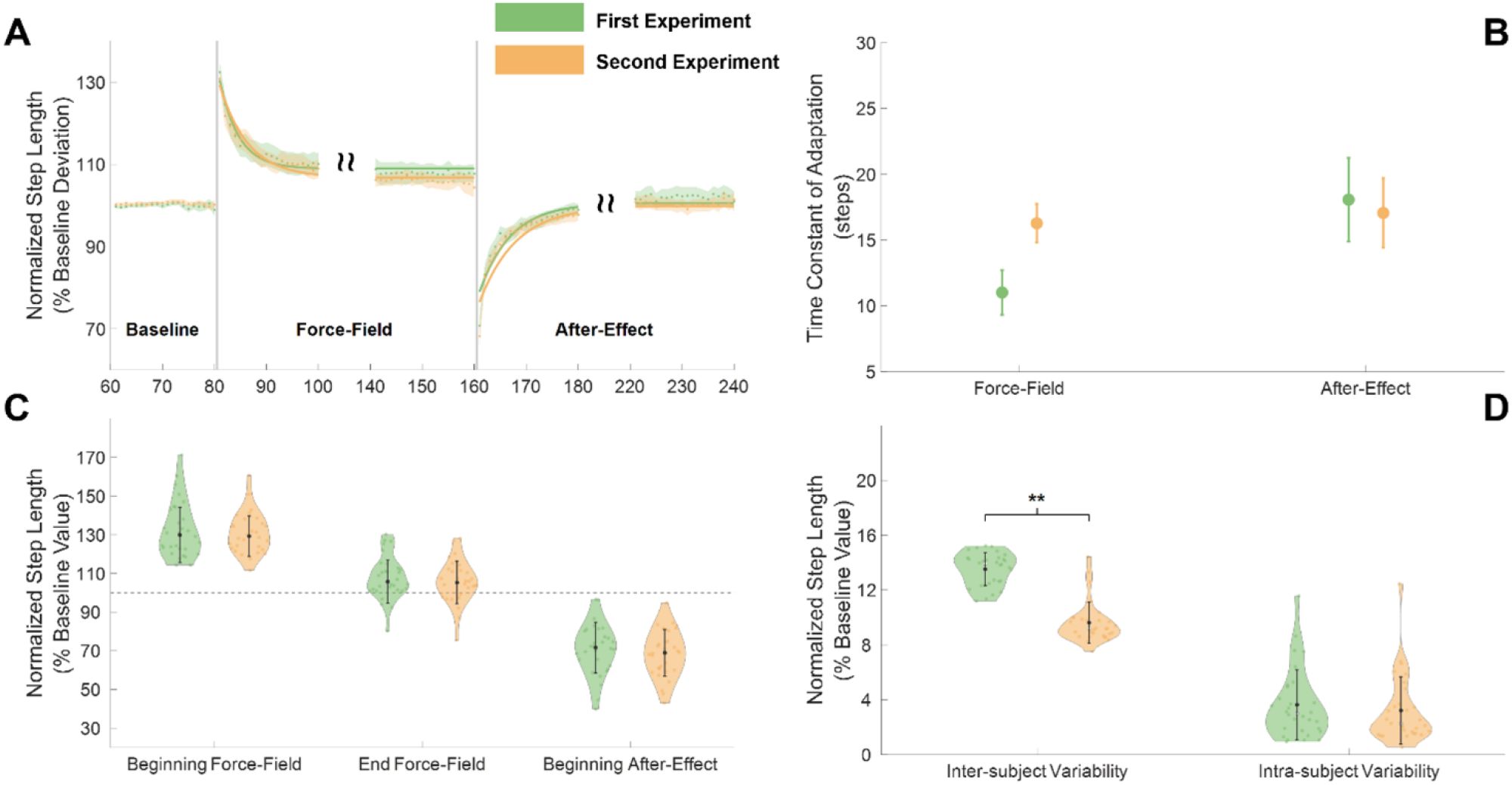
Analysis of step length adaptation, FIRST vs SECOND experiment, regardless of stimulation condition. (**A**) Step length changes over the course of the experiment for the FIRST (light green) and SECOND (yellow) experiments. Dots and shaded areas represent average and standard error of the normalized step length across all participants for each condition. Bold lines represent the exponential fitting calculated from the average data of each condition. (**B**) Time constants (in steps ± 95% confidence interval) of the exponentials estimated during the Force-field and After-Effect phases for the FIRST and SECOND experiments regardless of condition. (**C**) Across-participants average ± standard deviation of step length at the beginning and end of the Force-Field and at the beginning of the After-Effect. Error bars represent median ± standard deviation across participants. (**D**) Inter-subject and intra-subject variability in step length. Error bars represent median ± standard deviation across participants.

**Table S1.** Statistical analysis of notable points in the kinematic and kinetic plots (identified from Figure 4, original manuscript). All statistical analyses are based on Wilcoxon’s signed rank test.

|  | STIM | NOSTIM | p-value |
| --- | --- | --- | --- |
| Position peak hip flexion during swing (% Gait Cycle) | 84.67 ± 6.2 | 86.54 ± 6.5 | <0.01 |
| Position peak knee flexion during swing (%) | 69.52 ± 4.3 | 70.35 ± 4.2 | ns |
| Position peak knee extension during swing (%) | 95.83 ± 3.2 | 96.17 ± 4.1 | ns |
| Peak hip torque during swing (Nm/Kg) | 0.13 ± 0.03 | 0.13 ± 0.04 | ns |
| Position of peak hip torque during swing (%) | 88.27 ± 8.2 | 88.62 ± 7.8 | ns |
| Peak knee torque during swing (Nm/Kg) | 0.14 ± 0.05 | 0.14 ± 0.04 | ns |
| Position of peak knee torque during swing (%) | 78.93 ± 4.5 | 79.89 ± 5.1 | ns |
| Minimum knee torque during swing (Nm/Kg) | -0.07 ± 0.03 | -0.07 ± 0.04 | ns |
| Position of minimum knee torque during swing (%) | 91.91 ± 6.7 | 87.37 ± 12.12 | ns |
| Position of minimum hip power during late swing (%) | 90.38 ± 4.8 | 91.31 ± 5.0 | <0.05 |
| Peak knee power during late swing (W/Kg) | 0.84 ± 0.43 | 0.81 ± 0.49 | ns |
| Position of peak knee power during late swing (%) | 87.93 ± 7.6 | 87.81 ± 6.7 | ns |

